# Using stochastic dynamic programming for making decisions about invasive species

**DOI:** 10.64898/2026.09.05.749566

**Authors:** Olivier Gimenez, Abigail G. Keller, Lucile Marescot, Carl Boettiger, Cassie Speakman

## Abstract

1. Invasive species pose significant ecological, economic, and public health challenges. Their management requires repeated decisions under uncertainty about when, where and how intensively to intervene. Although structured decision-making and adaptive management provide conceptual frameworks for addressing these challenges, their practical implementation remains limited because optimization methods such as stochastic dynamic programming (SDP) are often perceived as mathematically complex, challenting to apply and difficult to interpret.
2. Here, we provide a practical guide to using SDP and its extensions for invasive species management. Using coypu (*Myocastor coypus*) management as a running case study, we introduce the framework through a minimal working example before progressively extending it to spatial dynamics, imperfect detection using partially observable Markov decision processes (POMDPs), adaptive management under model uncertainty, and value-of-information analyses that quantify the benefits of learning before acting.
3. All examples are accompanied by reproducible implementations in R, together with recommendations on model formulation, parameterization and interpretation. We also discuss the strengths and limitations of SDP.
4. By lowering the technical barrier to SDP, this guide aims to help ecologists and wildlife managers integrate decision theory into invasive species management, enabling transparent, reproducible and objective-driven management strategies under uncertainty. More broadly, the methods presented here are applicable to a wide range of ecological decision problems beyond invasive species, wherever management requires balancing immediate actions against uncertain future outcomes.

## 1. Introduction

Invasive species are a major global concern, with wide-ranging impacts on ecosystems, economies, and public health (Pyšek et al. 2020, Roy et al. 2024). Their management is challenging, as it requires repeated decisions under uncertainty, limited budgets, and imperfect knowledge of ecological dynamics, which are often characterized by rapid spread, strong population fluctuations, and broad spatial distributions. In practice, managers must decide not only whether to intervene, but also when, where, and how intensively, while balancing ecological, economic, and social consequences that may unfold over different time horizons (Hauser and Possingham 2008).

Structured approaches are therefore necessary for addressing the fundamentally dynamic nature of invasive species decision problems. Predicting invasion dynamics or mapping risks is not sufficient unless these predictions are explicitly linked to management objectives, feasible actions, and measurable consequences. Decision-making is further complicated by the need to account for multiple, sometimes competing, criteria, including ecological effectiveness, economic costs, operational feasibility, and potential unintended consequences such as non-target impacts. In addition, management actions are embedded in social contexts, where stakeholder perceptions, acceptability of control measures, and underlying environmental values can strongly influence both implementation and outcomes (Simberloff et al. 2013, Estévez et al. 2015, Crowley et al. 2017)

Structured decision-making and adaptive management have long been advocated as frameworks to address the complexity of ecological management problems (Williams et al. 2002, Thompson et al. 2021). These approaches emphasize the explicit formulation of objectives, the evaluation of alternative actions, and the incorporation of uncertainty into decision processes. However, despite their conceptual appeal, they remain underused in practice, in part because of the technical challenge of formally identifying and quantifying competing objectives reflecting different stakeholder values and priorities, and optimizing the resulting trade-offs over time.

Stochastic dynamic programming (SDP), a core tool of decision theory, provides a powerful framework to address this challenge by treating management as a sequential decision problem under uncertainty. In this framework, optimal actions are derived as a function of the current state of the system (e.g. population size or distribution across management units), while explicitly accounting for stochastic dynamics and the future consequences of present decisions. SDP naturally accommodates a wide range of management actions, including prevention, containment, control, eradication, surveillance, or inaction, and allows their outcomes to be evaluated in terms of long-term ecological, economic, or social objectives (Marescot et al. 2013).

SDP has been successfully applied to a variety of invasive species management problems, including optimizing prevention versus control strategies (Leung et al. 2002), designing control policies for invasive plants (Hyder et al. 2008), identifying state-dependent strategies for spatially structured invasions (Bogich et al. 2008), optimizing search effort for invasive insects (Baxter and Possingham 2011), and comparing alternative strategies across invasion stages (Hyytiäinen et al. 2013). The SDP framework can also be extended to imperfectly known system states, allowing monitoring and control of invasive populations to be jointly optimized (Waring et al. 2024).

Despite these advances, the practical implementation of SDP and related approaches remains limited in applied ecology. This gap is partly due to the perceived complexity of these methods and the lack of accessible, step-by-step guidance for translating ecological problems into operational decision models.

In this paper, we aim to bridge this gap by providing a practical guide to the use of SDP and its extensions for invasive species management. Using the coypu (*Myocastor coypus*) as a running example, we illustrate how to formulate decision problems, implement models, and derive optimal management strategies across a range of ecological contexts, from single populations to metapopulations. We first introduce the core concepts of SDP through a minimal, fully transparent example. We then develop a series of worked tutorials of increasing complexity, including spatial dynamics, imperfect detection, adaptive management under model uncertainty, and value of information analyses that quantify the benefits of learning before acting. Throughout, we provide reproducible implementations in R to facilitate uptake and adaptation by practitioners.

## 2. Explanation of the method

Here, we present a minimal SDP example. The aim is purely pedagogical: to get familiar with the terminology and understand how states, actions, transition probabilities, rewards, discount and time horizon combine to produce an optimal decision rule.

We consider a simplified invasion management problem with three qualitative states describing the invasion level: Eradicated with no individuals detected but possible re-invasion risk; Contained with the invasive species present but at low abundance; or Established with the invasive species widespread with high impacts.

At each time step *t*, the manager chooses between two actions: Monitor with surveillance only (low cost, little effect on the population); or Control with active management campaign (higher cost, greater effect).

Transition matrices describe how the invasion state evolves from one time step to the next under each management action. We denote action-dependent transition matrices *P_a_*(*s*, *s*^′^) where *a* denotes an action, *s* and *s*^′^ are for current and future states:

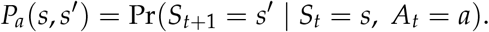

Each row corresponds to the current state and each column to the next state, with entries giving transition probabilities. We now build a transition matrix for each action. Under monitoring, the transition is:

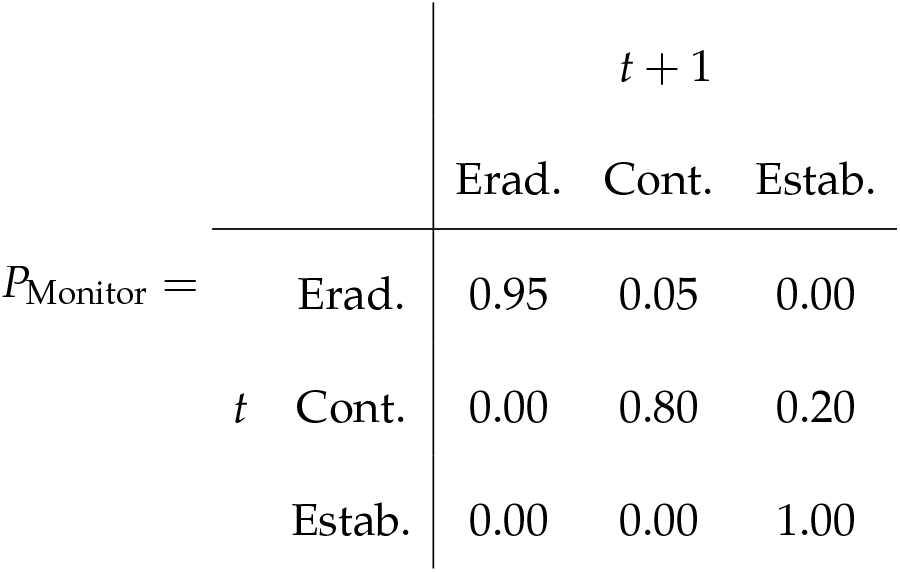

If the system is eradicated at *t*, it remains so at *t* + 1 with probability 0.95 but may revert to a contained state with probability 0.05, reflecting a small risk of re-invasion. When the invasion is contained, it persists with probability 0.80 but may deteriorate to an established state with probability 0.20, indicating possible population growth without intervention. Once established, the system remains established with probability 1, meaning monitoring alone cannot reverse the invasion.

In contrast, control shifts the system toward better states:

| | | $t + 1$ | | |
| --- | --- | --- | --- | --- |
|  |  | Erad. | Cont. | Estab. |
| $P_{\text{Control}} =$ | Erad. | 0.98 | 0.02 | 0.00 |
| | $t$ Cont. | 0.20 | 0.75 | 0.05 |
|  | Estab. | 0.05 | 0.75 | 0.20 |

From an eradicated state at *t*, the system remains eradicated at *t* + 1 with probability 0.98 and only rarely becomes contained (0.02), reflecting the challenges of complete eradication. When contained, control can eradicate the population (0.20), maintain it at low levels (0.75), or fail and allow establishment (0.05). Finally, when the invasion is established, control improves the situation in most cases, leading to a contained state with probability 0.75, occasionally achieving eradication (0.05), but sometimes failing to reduce impacts (remaining established with probability 0.20).

Together, these transition matrices encode both ecological uncertainty and the imperfect effectiveness of management actions.

Next we define the reward function denoted as *R*(*s*, *a*). Here, rewards are defined as the negative of total costs, so that maximizing expected reward is equivalent to minimizing management and damage costs:

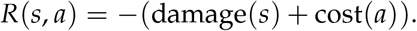

Rows correspond to the current invasion state and columns to the management action. Damage depends on the invasion state (higher when the invasion is established), while management cost depends on the chosen action (higher for active control than monitoring).

For this illustrative example, we assign arbitrary, unitless values to damage and management costs, chosen solely to generate a simple trade-off between ecological impacts and management effort. The reward matrix is then constructed by combining these two components for all state-action pairs:

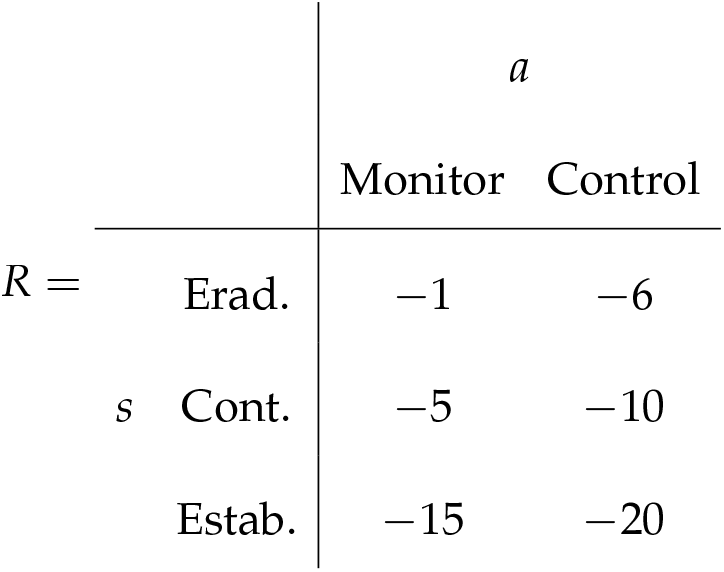

For example, when the invasion is eradicated, monitoring yields a reward of-1, representing relatively low surveillance costs, whereas applying control in the same state yields-6 due to higher intervention costs. When the population is contained, rewards are-5 for monitoring and-10 for control, representing moderate damages combined with the respective management costs. Finally, when the invasion is established, rewards drop to-15 under monitoring and-20 under control, reflecting both high ecological impacts and, in the latter case, substantial intervention effort. Overall, this reward structure captures the short-term trade-off between ecological impacts and management effort that underlies the decision problem.

Together, the transition matrices and reward function provide all the ingredients needed to formulate the SDP problem. The key idea is that management is a sequential decision problem: decisions are made repeatedly over time, and successive decisions are linked because an action taken at one time step affects the state of the system, and therefore the management options and outcomes, at subsequent time steps. The value of a management decision thus depends not only on its immediate consequences, but also on how it influences future states and opportunities. This principle is formalized by the Bellman equation (Bellman 2003),

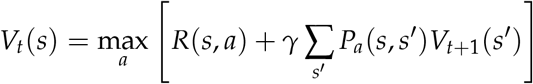

where *V_t_*(*s*), the value function, is the optimal expected value from state *s* at time *t* onward, and *γ* is the discount factor describing the relative importance of short-and long-term rewards. The value function *V_t_*(*s*) therefore summarizes the best outcome that can be expected from the current state, accounting for both the immediate reward and all future rewards obtained by following the optimal sequence of actions (Figure 1).

**Figure 1:**
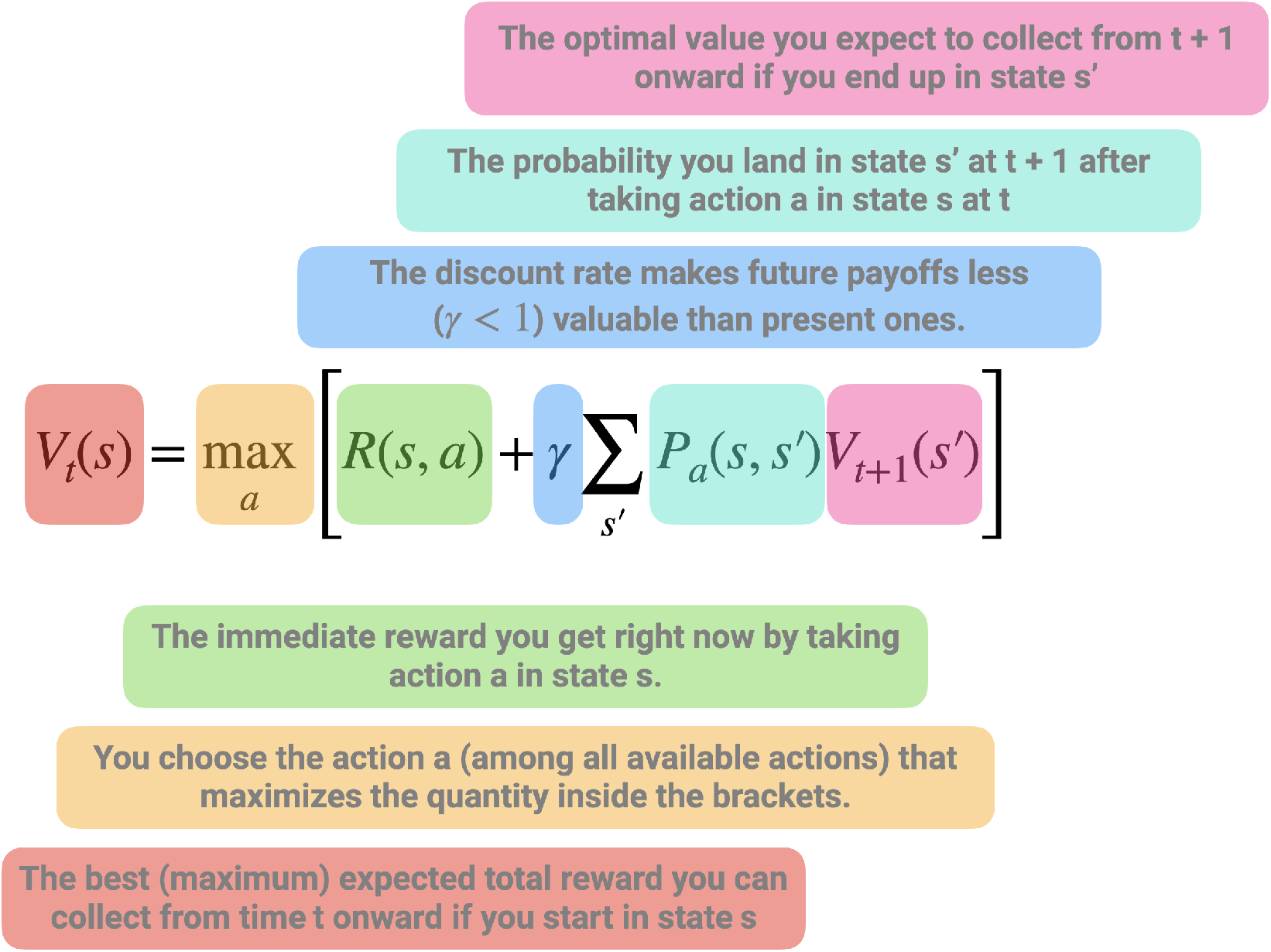
Representation of the Bellman equation in stochastic dynamic programming. The value of being in a given state at time *t*, *V_t_*(*s*), is defined as the maximum, over all possible actions, of two components: (i) the immediate reward obtained by taking action *a* in state *s*, and (ii) the expected discounted future value of subsequent states. The latter corresponds to the average (weighted by transition probabilities) of the optimal future values *V_t_*_+1_(*s*^′^) that can be reached from the current state-action pair, scaled by a discount factor *γ <* 1 to reflect that future payoffs are less valuable than present ones. This equation highlights the fact that the value of a decision is not only determined by its immediate payoff, but also by how it shapes future opportunities. The optimal action is therefore the one that maximizes the sum of current rewards and expected future benefits.

In this first example, we consider a finite-horizon decision problem, where management decisions are made over a fixed number of time steps, *t* = 1,…,*T*, with *T* = 5 in our example. Because no decisions remain after time *T*, we set the terminal value at *T* + 1 to *V_T_*_+1_(*s*) = 0 for every state. The optimal policy is then obtained by working backwards through time, a procedure known as backward induction, or backward iteration (Marescot et al. 2013). Thus, starting from the final time step *t* = *T*, the algorithm successively moves backwards through time to *t* = 1. At each time step and for each state, it evaluates every available management action, combines its immediate reward with the expected value of future states according to the Bellman equation, and retains the action that maximizes the total expected reward (Figure 1).

Applying the Bellman equation yields the value function *V_t_*(*s*), which represents the maximum expected cumulative reward (equivalently, the minimum expected cumulative cost) obtained when starting from state *s* at time *t* and following the optimal policy thereafter:

| | | $t = 1$ | $t = 2$ | $t = 3$ | $t = 4$ | $t = 5$ | $t = 6$ |
| --- | --- | --- | --- | --- | --- | --- | --- |
| $V_t(s) =$ | $s = \text{Erad.}$ | -7.60 | -5.50 | -3.69 | -2.20 | -1 | 0 |
| | $s = \text{Cont.}$ | -33.78 | -27.43 | -19.96 | -12.0 | -5 | 0 |
| | $s = \text{Estab.}$ | -49.26 | -42.05 | -34.47 | -26.8 | -15 | 0 |

Values are more negative when the invasion is established because higher ecological damages are expected to accumulate over time. Conversely, values become progressively less negative as the planning horizon shortens, reflecting the fact that fewer future costs remain to be incurred.

The associated optimal policy *π_t_*(*s*) specifies the best action to take in each state *s* at each time step *t*:

| | | $t = 1$ | $t = 2$ | $t = 3$ | $t = 4$ | $t = 5$ |
| --- | --- | --- | --- | --- | --- | --- |
| $\pi_t(s) =$ | $s = \text{Erad.}$ | Monitor | Monitor | Monitor | Monitor | Monitor |
| | $s = \text{Cont.}$ | Control | Control | Monitor | Monitor | Monitor |
| | $s = \text{Estab.}$ | Control | Control | Control | Control | Monitor |

The optimal policy illustrates one of the defining features of SDP: the best decision depends on the current state of the system. When the invasion has been eradicated, additional control provides little benefit relative to its cost, making monitoring the preferred strategy throughout the planning horizon. In contrast, once the invasion is established, immediate control is generally optimal because delaying intervention substantially increases expected future damages. The contained state lies between these two extremes, where the optimal decision depends on the remaining planning horizon and reflects the trade-off between the immediate cost of intervention and the long-term benefits of preventing further spread.

## 3. Things to consider for before using this method

Applying SDP to ecological problems requires careful formulation of the decision context, as well as practical considerations regarding model structure, data availability, and computational complexity.

### 3.1 Problem formulation

A prerequisite for SDP is a clear definition of the decision problem. This can be clarified through processes like structured decision making or the PrOACT framework, which includes specifying management objectives, identifying feasible actions, defining the state variables that describe the system, and constructing a reward function that captures the trade-offs among ecological, economic, and social outcomes (Marescot et al. 2013, Hemming et al. 2022). In the following examples, we assume pre-specified management objectives and available actions. Because SDP optimizes decisions given these elements, poorly specified objectives or unrealistic action sets can lead to misleading recommendations (Gregory et al. 2012, Runge et al. 2020).

### 3.2 State definition and discretization

SDP typically requires a finite state space, whereas ecological systems are often described by continuous variables (e.g. abundance, spatial extent). Discretization is therefore necessary, but introduces a trade-off between realism and tractability. Fine discretization improves accuracy but increases computational cost, while coarse discretization may obscure important system dynamics (Nicol and Chadès 2012, Chadès et al. 2021). This trade-off becomes increasingly important as the number of state variables and management actions increases, leading to the curse of dimensionality (Walters and Hilborn 1978). In practice, sensitivity analyses can be used to assess whether management recommendations are robust to the chosen discretization. When computational costs become prohibitive, possible solutions include simplifying the model, aggregating states, or using simulation-based approaches to approximate optimal policies (Nicol and Chadès 2011, Schapaugh and Tyre 2012, Marescot et al. 2013).

### 3.3 Model specification and uncertainty

The performance of SDP depends critically on the specification of transition dynamics and reward functions, which are often uncertain in ecological systems. In practice, managers rarely know the true probabilities governing population dynamics, the effectiveness of alternative management actions, or the ecological and economic consequences associated with different system states (Polasky et al. 2011). These uncertainties may arise from limited data, structural assumptions, or imperfect knowledge of ecological processes (Regan et al. 2002). Ignoring them can lead to suboptimal management recommendations, motivating extensions such as partially observable Markov decision processes, which explicitly account for imperfect observations, and adaptive management, which incorporates learning about uncertain system dynamics over time (Haight and Polasky 2010, Williams et al. 2011, Chadès et al. 2021).

### 3.4 Data requirements and interpretation

Although SDP can be implemented using simple or expert-based models (Moore and McCarthy 2016, Chenery et al. 2020), its usefulness ultimately depends on the quality of the ecological understanding used to parameterize the decision model. In adaptive management applications, learning is only possible if the candidate model set adequately represents the true system dynamics (Runge et al. 2016). If the true dynamics lie outside the range of considered hypotheses, or if ecological conditions change in unexpected ways (e.g. under climate change), management recommendations may become unreliable despite continued monitoring and learning (Tucker and Runge 2021, Keller et al. 2025). In data-limited contexts, SDP may still provide valuable qualitative insights into decision structure and trade-offs, but quantitative recommendations should be interpreted with caution (Baxter et al. 2007).

## 4. Worked examples

Throughout this paper, we use the coypu as a running example to illustrate SDP. Native to South America, the coypu was introduced into Europe during the early twentieth century through the fur trade and has since established widespread invasive populations across many European countries (Bonnet et al. 2023). Coypus damage crops, riverbanks and hydraulic infrastructures, while also acting as reservoirs of zoonotic pathogens such as Leptospira spp., making them both an ecological and public health concern (Diagne et al. 2023, Bonnet et al. 2023). Their widespread distribution, rapid population growth and the need for repeated control interventions make coypus an ideal case study for illustrating sequential decision-making under uncertainty.

The worked examples presented here build on the coypu case study developed in Gimenez (2025), extending it from estimating population abundance to optimizing management decisions. The examples address realistic invasive species management problems, but the decision models and parameter values were developed for pedagogical purposes rather than for direct operational implementation. Their formulation nevertheless draws on our experience with invasive rodent management and reflects plausible ecological dynamics, management actions, objectives, and trade-offs encountered in these systems.

In what follows, we use the R package MDPtoolbox v4.0.3 (Chadès et al. 2014) to compute optimal policies using SDP, unless stated otherwise.

### 4.1 Regulating a single invasive population

The objective of this example is to determine an optimal regulation policy for coypu based on their total abundance. We consider a management problem in which a decision maker chooses, at each time step, the intensity of a control effort *u* ∈ [0, 1]. Low effort corresponds to minimal intervention, while high effort represents intensive removal. The management objective is to balance three competing components: ecological damages that increase with population size (e.g. crop losses, dike weakening), management costs that increase with effort, and an additional penalty if abundance exceeds a threshold level.

In contrast to the previous example with discrete invasion states, the system is here described by a continuous state variable, the population size *N*. Because dynamic programming requires a finite state space, we discretize abundance onto a grid ranging from 0 to the carrying capacity *K*. In the implementation, we set *K* = 2000 individuals and use a grid step of 20 individuals. Thus, we have a total of 100 states representing the abundance of coypu. Control effort is also discretized, from 0 to 1 in increments of 0.05 (i.e. 20 actions).

Population dynamics follow a density-dependent growth model with removals due to control,

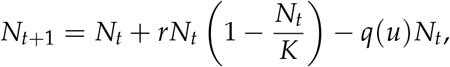

where the first term describes logistic growth and *q*(*u*) is the fraction of individuals removed as a function of effort. We use a saturating removal function,

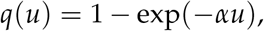

with *α* = 1.5, which implies diminishing returns of control effort (Figure 2A). For example, an effort of *u* = 0.5 removes about 53% of the population, whereas maximum effort *u* = 1 removes about 78%, reflecting the fact that complete eradication is rarely achievable in a single step.

**Figure 2:**
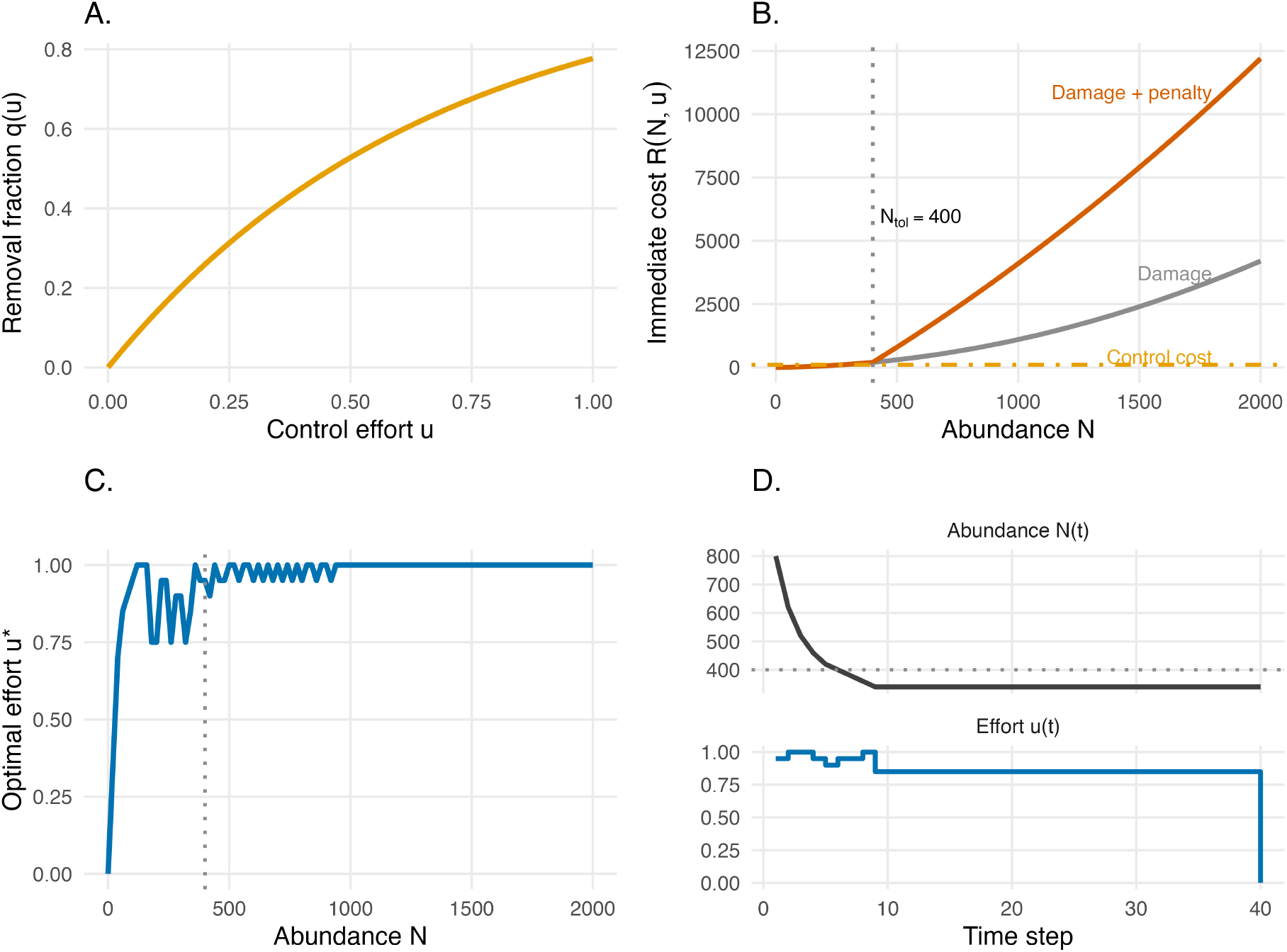
Components and outputs of an infinite-horizon stochastic dynamic programming model for invasive species management. (A) Removal fraction *q*(*u*) as a function of control effort *u*. (B) Immediate cost components as functions of abundance. Damage increases nonlinearly with population size, and above the tolerable threshold (*N*_tol_ = 400), an additional penalty is added, resulting in rapidly increasing total costs. Control cost is shown for an illustrative effort level (*u* = 0.6). (C) Optimal stationary policy *u*^∗^(*N*), showing the recommended control effort for each abundance level; the dotted vertical line indicates the tolerable abundance threshold. (D) Example trajectories of abundance and optimal control effort under the policy, starting from *N*_0_ = 800; the dotted horizontal line indicates *N*_tol_.

Rewards are defined as the negative of total costs, so that maximizing expected reward is equivalent to minimizing long-term damages and management expenses (Figure 2B):

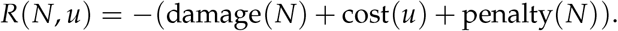

Damages increase non-linearly with abundance, using a simple quadratic form damage(*N*) = 0.1*N* + 0.001*N*^2^, reflecting the common ecological pattern whereby impacts accelerate at high density. Management costs increase with effort and are assumed convex (cost(*u*) = 50*u* + 200*u*^2^), capturing increasing marginal costs of intensive interventions. Finally, a penalty is applied when abundance exceeds a tolerable threshold *N*_tol_ = 400, representing situations where ecological or social impacts become unacceptable. The penalty increases linearly beyond this threshold according to

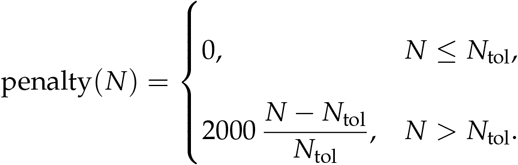

This formulation implies that the penalty is zero below the threshold and then rises proportionally to the exceedance. For example, an abundance of *N* = 800 yields a penalty of 2000, thereby strongly incentivizing control once populations grow beyond acceptable levels.

Unlike the previous toy example, we no longer assume a fixed planning horizon. Instead, management is assumed to continue indefinitely. In this setting, the decision problem is solved using infinite-horizon discounted value iteration (discount factor *γ* = 0.99), an iterative algorithm that repeatedly applies the Bellman equation until the value function converges (Marescot et al. 2013). The algorithm returns both a stationary optimal policy, which maps each abundance class to the recommended control effort, and a value function summarizing the expected long-term net benefit of following this policy from any initial abundance. Because the planning horizon is infinite, the optimal decision depends only on the current abundance and not on time.

The optimal policy exhibits a threshold-like pattern (Figure 2C). Control effort remains low when abundance is low because ecological damages are limited and additional removals provide little long-term benefit. As abundance approaches the tolerable threshold (*N*_tol_ = 400), effort increases rapidly, reflecting the growing ecological and economic consequences of delaying intervention. Above this threshold, intensive control becomes optimal to prevent sustained high damages and penalties. More generally, this result illustrates a central feature of SDP: optimal management decisions emerge naturally from balancing immediate intervention costs against expected future consequences.

Figure 2D illustrates the resulting population dynamics for a system initially containing *N*_0_ = 800 individuals. At each time step, the control effort is adjusted according to the current abundance, and the population responds through the logistic growth model with removals. In this deterministic example, abundance rapidly declines from its initial level before converging towards an equilibrium where the marginal benefit of additional control is balanced by its marginal cost.

### 4.2 Coordinating control across connected populations

We now extend the previous example from a single population to a spatially structured invasion in which local populations are connected through dispersal. The aim is to illustrate how SDP can coordinate management across multiple sites when local extinction and recolonization interact. We consider a network of eight sites, representing cities or management units, connected by an illustrative dispersal network. The observed coypu abundances reported in Gimenez (2025) are used only to define the initial pattern of occupied and unoccupied sites; thereafter, the decision model operates directly on occupancy states.

At time *t*, each site *i* is either occupied or empty,

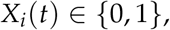

and the state of the whole system is the vector

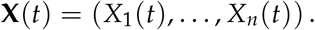

With *n* = 8 sites, there are 2^8^ = 256 possible spatial configurations. This is an important distinction from the previous worked example: rather than using local abundance as the state variable, we focus here on the spatial extent and configuration of the invasion.

At each time step, the manager selects one of three global management actions which is applied across the network: no, moderate, or strong control. Management affects two components of the invasion process. First, occupied sites may become locally extinct. In the absence of control, local extinction is unlikely, whereas increasingly intensive control raises the probability that an occupied site becomes empty (Figure 3A). We denote these action-dependent probabilities by

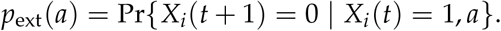

**Figure 3:**
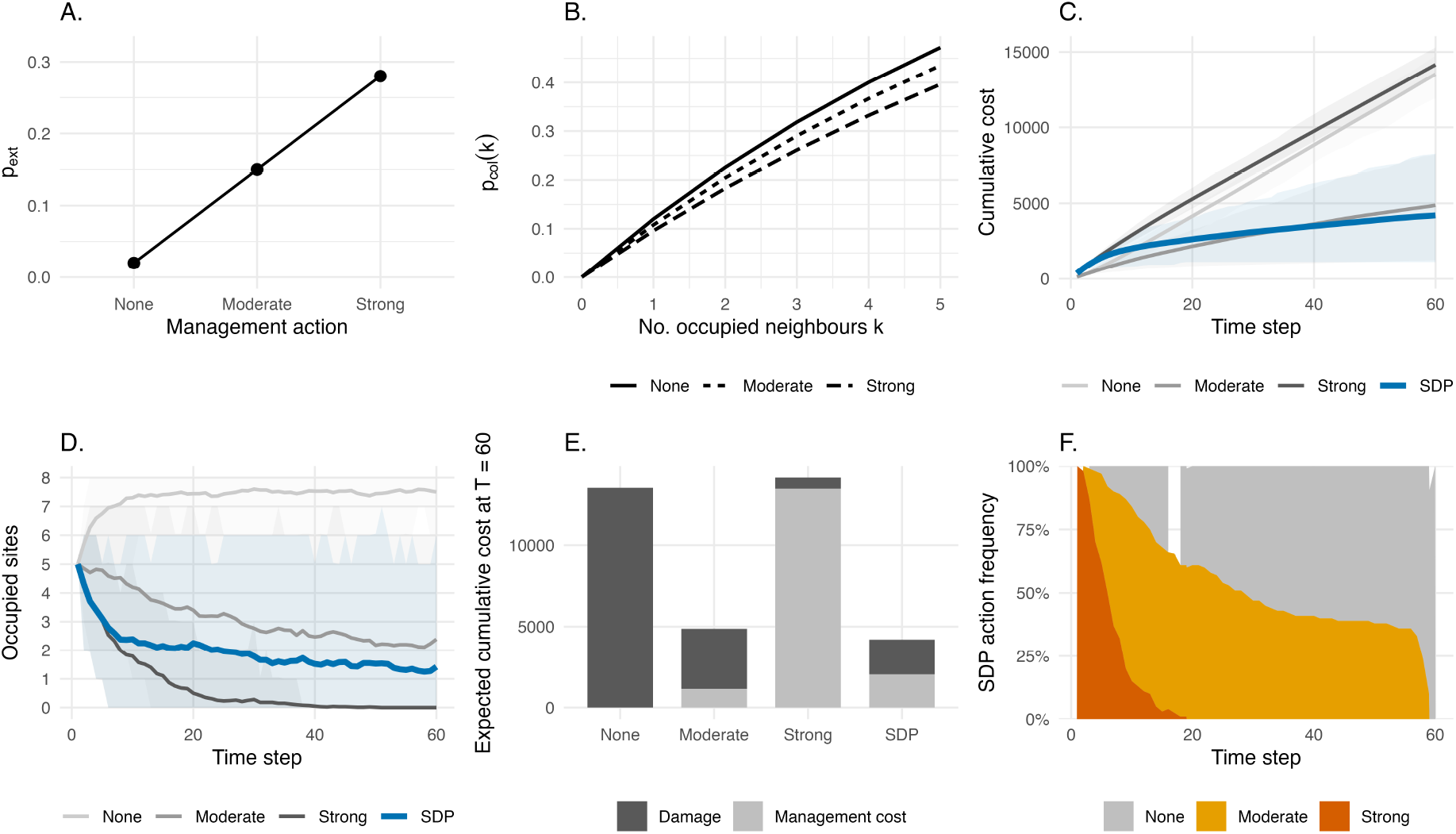
Spatial stochastic dynamic programming for coordinated management of connected invasive populations. (A) Local extinction probability under no, moderate, and strong control. (B) Colonization probability as a function of the number of occupied neighbouring sites under each management action. (C) Mean cumulative cost through time under three constant management strategies and the state-dependent SDP policy; shaded areas represent 90% simulation intervals. (D) Mean number of occupied sites through time under each strategy, with 90% simulation intervals. (E) Decomposition of expected cumulative cost at the end of the planning horizon into ecological damage and management expenditure. (F) Proportion of simulated trajectories in which the SDP selects no, moderate, or strong control at each time step.

In our illustrative example, these probabilities are set to 0.02, 0.15, and 0.28 under no, moderate, and strong control, respectively.

Second, empty sites may be recolonized from neighbouring occupied sites, as defined by the dispersal network. Let

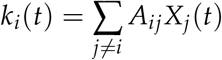

denote the number of occupied neighbours of site *i*, where *A_ij_*is the adjacency matrix describing connections among sites, with *A_ij_*= 1 if sites *i* and *j* are connected and *A_ij_* = 0 otherwise. Colonization probability increases with *k_i_*(*t*) according to

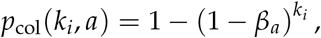

where *β_a_* is the effective per-neighbour colonization probability under management action *a*. Control is assumed to reduce colonization pressure, for example by lowering source populations or limiting dispersal from occupied sites. Consequently, recolonization becomes increasingly likely as more neighbouring sites are occupied, but this risk is reduced under stronger management (Figure 3B).

Together, local extinction and recolonization generate spatial feedbacks: management at one time step can influence not only whether individual sites remain occupied, but also the future risk of invasion elsewhere in the network. For each current spatial configuration and management action, we calculate the probability of every possible configuration at the next time step. Although the colonization probability of a site depends on the occupancy state of its neighbors, this dependence is fully determined by the current network configuration. We therefore assume that site-level transitions at the next time step are conditionally independent given the complete current network state and management action, allowing the transition probability between two network states to be obtained by combining the site-specific probabilities. Because the state space contains only 256 configurations, all states and transitions can be enumerated exactly. For larger or more complex state spaces, where exhaustive enumeration becomes computationally prohibitive, approximation methods such as heuristic sampling can instead be used to approximate optimal management policies (Nicol and Chadès 2011).

Management decisions must balance the ecological costs associated with a widespread invasion against the costs of intervention. We assume that damage increases non-linearly with the number of occupied sites,

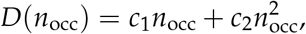

where *n*_occ_ is the current number of occupied sites. The quadratic component represents accelerating impacts as the invasion spreads across the network. Management costs contain both a fixed cost of implementing action *a* and a component that increases with the number of occupied sites,

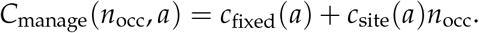

Strong control is therefore more effective at promoting local extinction and limiting recolonization, but it is also substantially more expensive to maintain. The immediate reward depends on the current spatial configuration (*s*) through its number of occupied sites (*n*_occ_), and is defined as the negative of total cost,

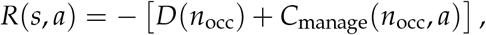

so that maximizing expected reward is equivalent to minimizing long-term ecological damage and management expenditure.

Here, the reward depends on spatial configuration only through the number of occupied sites. More generally, however, spatial structure can be incorporated directly into the reward function. For example, ecological damage could depend on which sites are occupied, their connectivity, or other landscape characteristics, while management costs and effectiveness could vary among sites or spatial configurations. This flexibility also allows objectives defined at different spatial scales to be combined within the same decision problem.

As in the introductory toy example, management is considered over a finite planning horizon, this time of 60 time steps. We therefore solve the SDP by backward induction, using a discount factor of *γ* = 0.99, providing almost equal weight between current and future rewards. The resulting policy specifies the optimal management action for every spatial configuration and time step. Importantly, two states containing the same number of occupied sites need not receive the same action because their spatial configurations may differ, as well as their future recolonization risks.

To evaluate the consequences of state-dependent management, we compared the SDP policy with three constant strategies in which no, moderate, or strong control is applied throughout the entire planning horizon. Because invasion dynamics are stochastic, each strategy was simulated repeatedly (100 times) rather than evaluated from a single trajectory. Figure 3C shows the resulting mean cumulative costs and their 90% simulation intervals. Applying strong control continuously can achieve substantial ecological benefits, but at a high management cost. In contrast, the SDP balances these competing objectives by changing intervention intensity as the state of the network changes, thereby reducing expected cumulative costs.

The same trade-off is apparent in the ecological outcomes (Figure 3D). Strategies differ in the expected number of occupied sites through time, with stronger control generally reducing invasion extent more rapidly. However, minimizing invasion extent alone is not necessarily optimal because maintaining strong control after the invasion has been substantially reduced can impose management costs that exceed the additional ecological benefits. Decomposing expected cumulative costs into ecological damage and management expenditure makes this trade-off explicit (Figure 3E).

Finally, Figure 3F shows the frequency with which the SDP selects each management action across simulated trajectories. Rather than prescribing a single intervention intensity throughout the study period, the optimal policy switches among control strategies as invasion risk changes. This illustrates the main benefit of SDP in a spatial management context: management intensity can be coordinated through time in response to the current configuration of the invasion, rather than applying the same level of control everywhere and indefinitely.

### 4.3 Managing under imperfect detection

The previous worked examples assumed that the manager always knows the true invasion state before making a decision. In practice, this assumption is rarely satisfied. Invasive species are often difficult to detect, particularly at low abundance or during the early stages of an invasion. Decisions must therefore be made under uncertainty, using imperfect monitoring information rather than the true ecological state. Partially observable Markov decision processes (POMDPs) provide a natural framework for addressing this problem by combining ecological dynamics with an explicit model of the observation process (Chadès et al. 2021, Waring et al. 2024). POMDPs are characterized by a tuple *< S*, *A*, *O*, *P*, *Z*, *r*, *γ*, *b*_0_ *>*, where *S*, *A*, *P*, *r* and *γ* are defined as before (see Section 2), *O* represents the set of observations *o* that a manager observes, *Z* is the observation function linking the observations to the states, and *b*_0_ is the initial belief state.

Again, we illustrate the approach using a coypu example. Here, we build upon the initial toy example where we have one population and a discrete state space. The ecological system can occupy one of three invasion states: Eradicated, Contained or Established, while the manager chooses one of three management actions: Do nothing, Fertility control or Lethal control.

The ecological dynamics are described by the set transition probabilities, similar to those used in the introductory example (see Section 2). Doing nothing allows contained populations to become established, fertility control partially reduces invasion pressure, and lethal control provides the highest probability of returning the system towards eradication, such that:

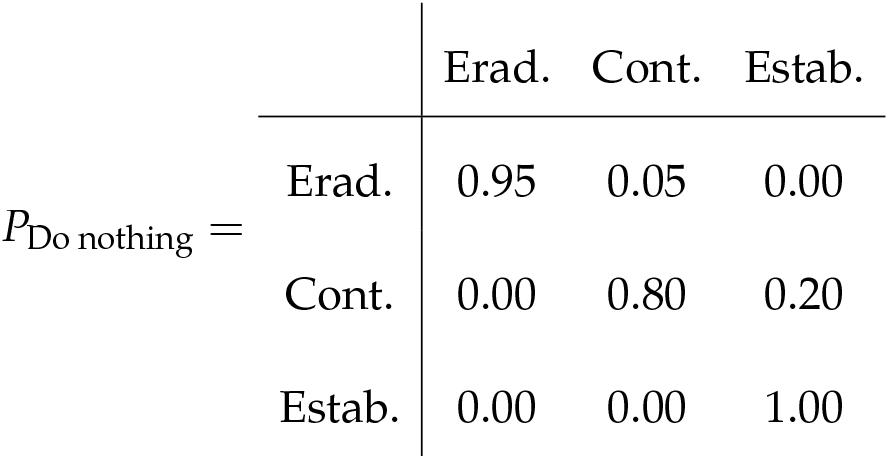
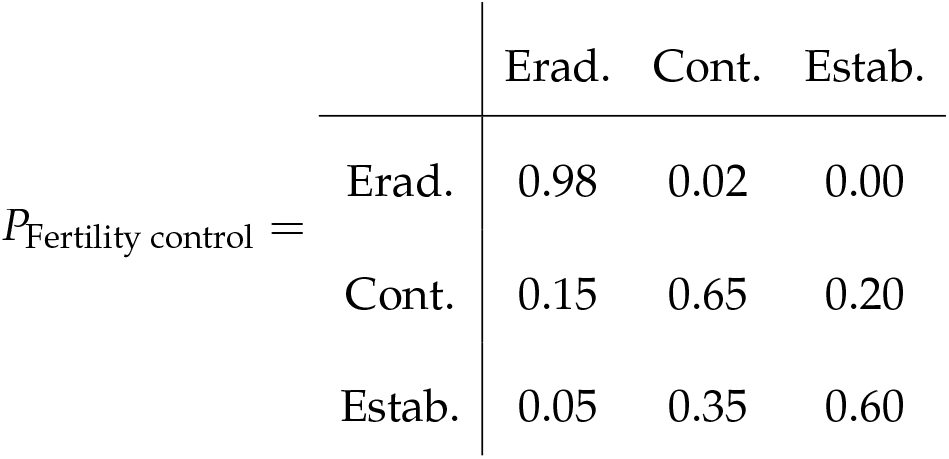
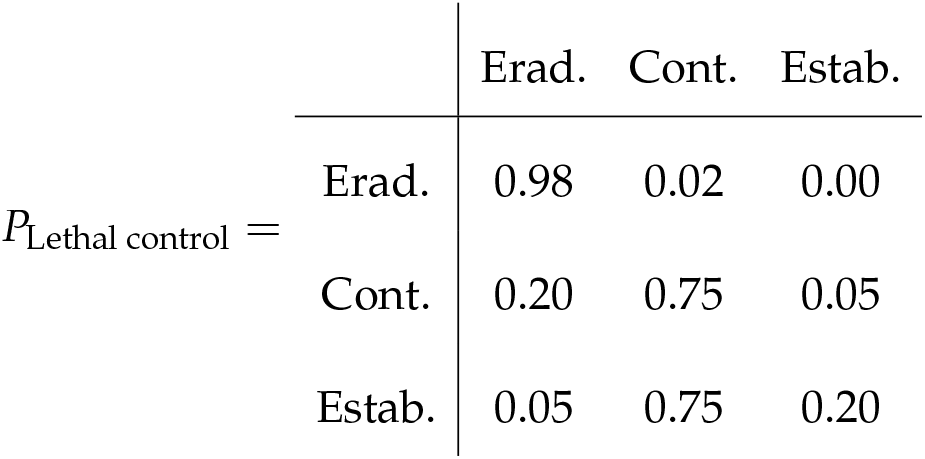

Unlike the previous examples, the invasion state is no longer directly observable. Instead, after each management action the manager receives an imperfect observation,

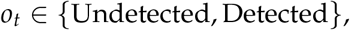

whose probability depends on both the true ecological state and the management action,

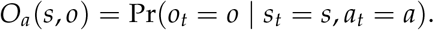

In this illustrative example, active management improves detection because field operations increase opportunities to observe coypu. Consequently, detection probabilities are higher under fertility or lethal control than when no management is undertaken (Figure 4A). Detection also increases with invasion severity, with established populations being more likely to be detected than contained populations.

**Figure 4:**
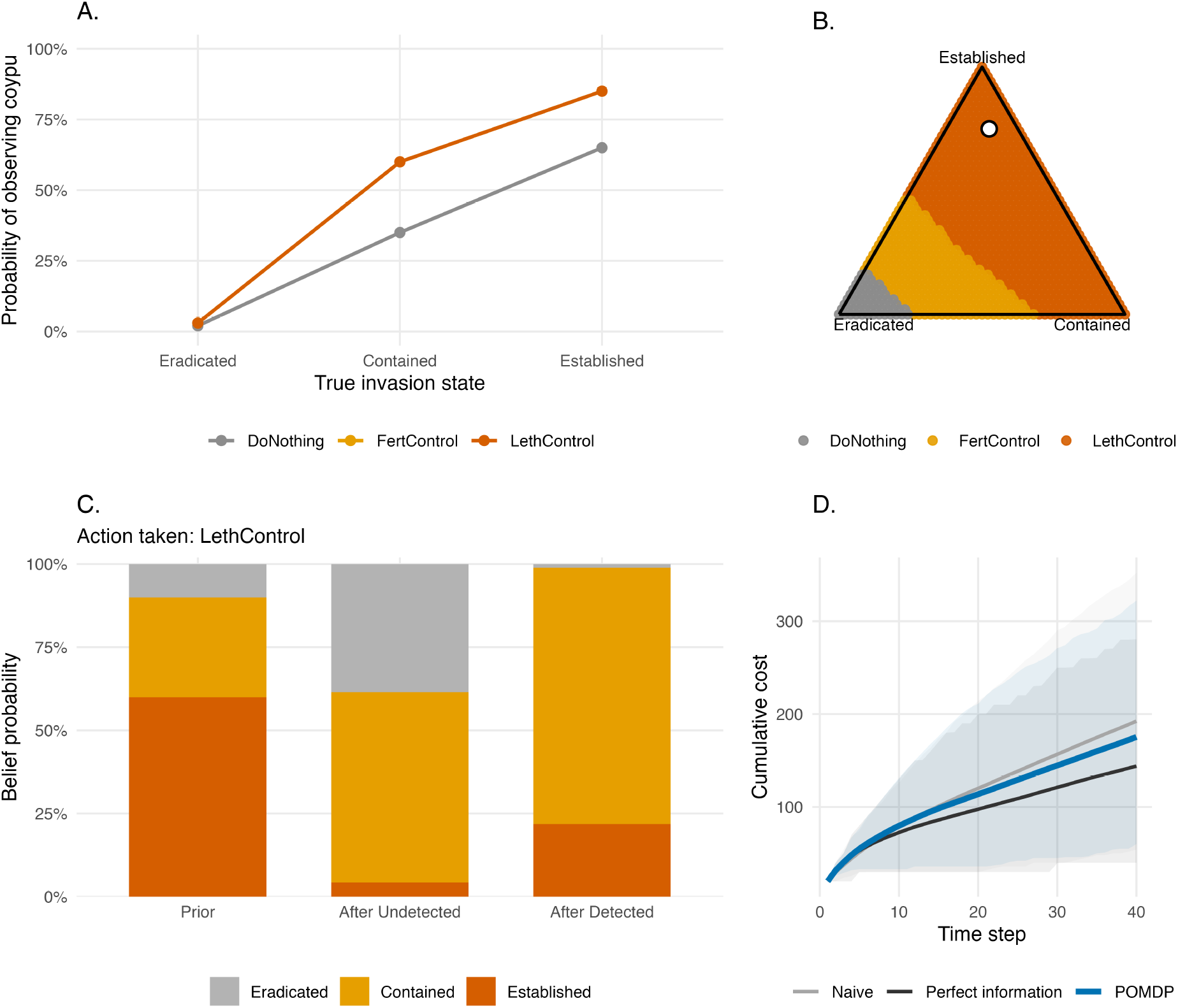
Managing invasive species under imperfect detection using a partially observable Markov decision process (POMDP). (A) Probability of detecting coypu as a function of the true invasion state under each management action. Active management is assumed to improve detection relative to no intervention. (B) Optimal management policy across the belief simplex. Each point represents a possible belief state (probability distribution over the three invasion states), and colours indicate the optimal management action. The white point corresponds to the example belief state used in the text (i.e., optimal action is lethal control if the probability of eradication, containment, and establishment are 0.10, 0.15, and 0.75, respectively). (C) Bayesian updating of the belief state following either a non-detection or a detection. The updated belief depends jointly on the previous belief, the ecological transition model and the observation model. (D) Mean cumulative management cost through time under three strategies: a naive observation-based policy, the optimal POMDP policy, and an ideal benchmark assuming perfect information. Shaded areas represent 90% simulation intervals.

Because the true invasion state is unknown, decisions are based on a belief state, denoted *b_t_*, rather than on the ecological state itself. The belief state is a probability distribution over all possible invasion states,

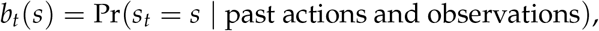

with

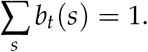

For example,

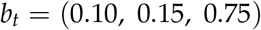

indicates that the manager believes that the system is most likely to be in an established state, while eradicated and contained are similarly probable. Geometrically, every possible belief corresponds to a point within the triangular belief simplex shown in Figure 4B. Rather than mapping ecological states to management actions, the optimal policy maps belief states to actions.

After each observation, the belief state is updated using Bayes’ rule by combining the previous belief, the ecological transition model and the observation model. Intuitively, the manager first predicts how the invasion is expected to evolve under the chosen management action and then revises this prediction after observing whether coypu are detected or undetected. Figure 4C illustrates this process for a representative belief state. Following a non-detection, the probability that the invasion has been eradicated increases substantially, whereas a detection shifts belief towards the contained and established states. These updated beliefs then determine the next management decision.

As in the previous examples, rewards are defined as the negative of total costs,

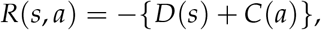

where ecological damage increases from eradicated to established states and management costs increase from doing nothing to lethal control. The POMDP is solved using the pomdp package (Hahsler and Cassandra 2025), which approximates the value function over the continuous belief space using a set of *α*-vectors. Each *α*-vector represents the expected long-term value of a particular management strategy for each possible ecological state; combining it with the current belief state gives the expected value of that strategy under the manager’s current uncertainty. The optimal action is then associated with the *α*-vector yielding the highest expected value.

To illustrate the benefit of explicitly accounting for imperfect detection, we compared the POMDP policy with two alternative strategies. The first is a naive strategy that bases decisions solely on the most recent observation (detected versus undetected), implicitly assuming that observations perfectly reveal the invasion state. The second is an idealized benchmark in which the true ecological state is known without error. Repeated forward simulations show that the POMDP consistently reduces long-term management costs relative to the naive strategy, although some performance is inevitably lost compared with the unrealistic case of perfect information (Figure 4D). This example highlights the central advantage of POMDPs: management decisions are informed not only by what has been observed, but also by the uncertainty associated with those observations.

### 4.4 Learning while managing under model uncertainty with adaptive management

So far, we assumed that the ecological model governing population dynamics was known. In reality, managers are often uncertain about the mechanisms driving population growth. Decision theoretic methods can be used to make control decisions not only in the face of uncertainty, but also to reduce uncertainty over time. Adaptive management, or “learning by doing”, involves making decisions to achieve a management objective while simultaneously learning about system dynamics to improve future management success (Holling 1978, Walters 1986, Conroy and Peterson 2013). Adaptive management approaches differ based on whether they learn only from the past (passive adaptive management) or anticipate future learning (active adaptive management) (Chadès et al. 2017).

Here we apply adaptive management strategies to inform control decisions for a single coypu population and to learn about and reduce uncertainty in the growth rate. We use the following discrete-time population model, where *r* is the intrinsic rate of growth; *K* is the carrying capacity; *q*(*u*) is the harvest rate as a function of action *u*; *R* is the per-capita growth rate; *m* describes whether the per-capita growth rate is linear, concave, or convex; and *ɛ* is log-normally distributed noise:

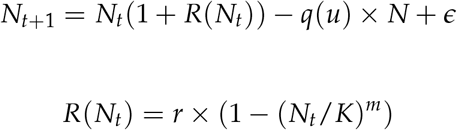

If *m >* 1, most density-dependent change occurs at high population levels (close to the carrying capacity), which is common for species with life history strategies typical of large mammals. If *m <* 1, most density-dependent change occurs at low population levels, which is common for species with life history strategies typical of insects and some fishes (Fowler 1981).

We consider a model set with five possible values of *m*. This model bounds, but does not contain, the true model we use to simulate dynamics in the passive adaptive management approach (Figure 5A). These different values of *m* result in different optimal policies when system dynamics are known perfectly (Figure 5B).

**Figure 5:**
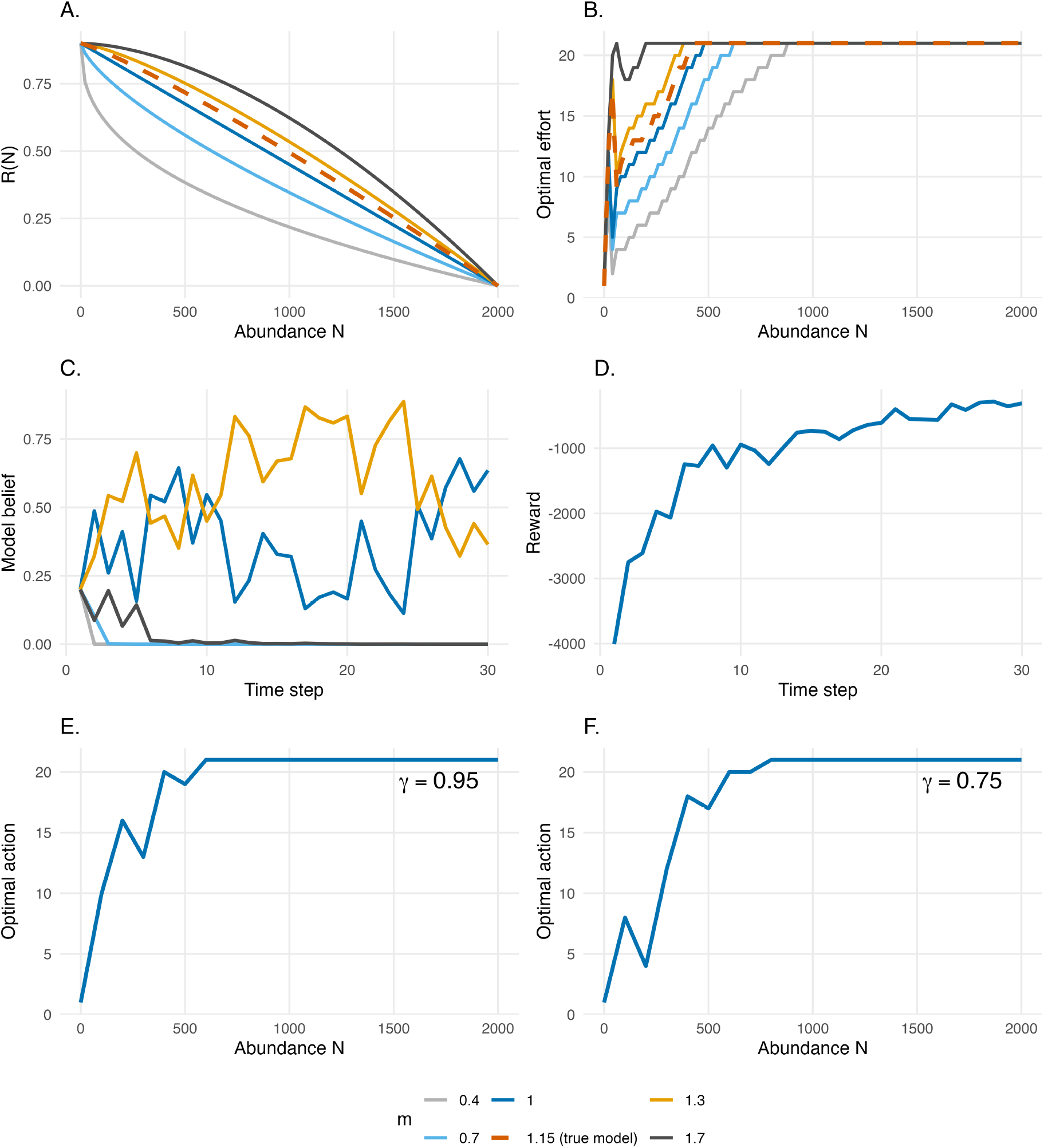
Adaptive management under structural uncertainty in population dynamics. (A) Model set representing structural uncertainty in the per-capita growth rate. (B) Optimal state-dependent removal policies assuming each model is known perfectly and solved using stochastic dynamic programming. Colours in (A-B) correspond to different values of *m*, with the red dashed line representing the true population dynamics used in the adaptive management examples. (C) Evolution of the belief assigned to each candidate population model through time under passive adaptive management; colours correspond to different values of *m*, representing alternative hypotheses about density dependence. (D) Reward obtained through time as management decisions are updated using the current belief state. As learning progresses, belief concentrates on the models closest to the true dynamics, leading to improved management performance. (E-F) Optimal state-dependent removal policies under active adaptive management for an initial uniform belief across models, with discount factors of (E) *γ* = 0.95 and (F) *γ* = 0.75. The higher discount factor places greater value on future rewards and hence on opportunities for learning, whereas the lower discount factor gives relatively greater weight to short-term rewards.

#### 4.4.1 Value of Information

For a decision maker, uncertainty may not be important to resolve; learning may not be inherently valuable if it does not result in improved management outcomes. A value of information (VoI) analysis a priori quantifies the importance of reducing uncertainty, or the expected improvement on management objectives if uncertainty were resolved (Raiffa and Schlaifer 1961). If the VoI is high, an adaptive management approach is typically justified.

The expected value of perfect information (EVPI) is measured as the difference between 1) *PI*, or the expected value once uncertainty has been resolved because the optimal action is chosen after knowing which model best describes the system, and 2) *NL*, or the expected value in the face of uncertainty because the optimal action is chosen with no learning about the system (Runge et al. 2011). The difference between these terms, EVPI = *PI* − *NL*, is the expected improvement in management performance due to acquiring perfect information. EVPI is expressed in the same units as the management objective: a value close to zero indicates that resolving uncertainty would have little effect on management performance, whereas a large EVPI indicates that reducing uncertainty could substantially improve outcomes. EVPI can also be interpreted as an upper bound on the resources worth investing to completely resolve the uncertainty.

To calculate EVPI for a sequential decision problem, we first use SDP to calculate the optimal state-dependent policy with assumed system dynamics, *π*^∗^(*s*)*_k_*, for each model *k* in the model set *K*, given each transition matrix, *P_k_* (Figure 5A-B). We also calculate the optimal policy given uncertain dynamics (or bet-hedging strategy), *π*^∗^(*s*)*_NL_*. We denote *b_t_*(*k*) as the belief assigned to model *k*, representing the current probability that model *k* adequately describes the system dynamics, with ∑*_k_ b_t_*(*k*) = 1. In the following example, we initially assign equal belief to all candidate models. This bet-hedging strategy is calculated with the transition matrix *P_NL_*, representing the average of the transitions of each model, *P_k_*, weighted by each model belief, *b_t_*:

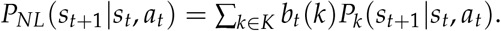

We calculate the EVPI through simulation of *i* replicated population trajectories using each model *k* in *K*. For every time step of the simulation over time horizon *T*, we apply both the optimal policy associated with model *k*, *π*^∗^(*s*)*_k_*, and the bet-hedging strategy, *π*^∗^(*s*)*_NL_*. The value of applying the optimal policy associated with the true dynamics is *V_k_*_,*t*,*i*_(*s_t_*, *a_t_*|*a_t_*= *π*^∗^(*s*)*_k_*), and the value of applying the bet-hedging strategy is *V_k_*_,*t*,*i*_(*s_t_*, *a_t_*|*a_t_*= *π*^∗^(*s*)*_NL_*). The EVPI is therefore *E_s_*_,*k*,*t*,*i*_[*V*(*s_t_*, *a_t_*|*a_t_* = *π*^∗^(*s*)*_k_*)] − *E_s_*_,*t*,*i*_[*V*(*s_t_*, *a_t_*|*a_t_* = *π*^∗^(*s*)*_NL_*)]. In other words, the EVPI is the average increase in value if the system dynamics were known perfectly and a manager acts optimally given those system dynamics, relative to the value if a manager makes the optimal decision without learning system dynamics.

In the context of coypu regulation, the EVPI represents the expected improvement on the management objective if uncertainty about the per-capita growth rate was reduced. We calculate the EVPI for a model set, *K*, where *m* ∈ {0.4, 0.7, 1, 1.3, 1.7} for *I* = 200 iterations over a *T* = 40 time horizon. We assume equal belief in each model. The expected value once uncertainty has been resolved, *PI*, is-1730.4, the expected value with no learning, *NL*, is-1787.8, and the EVPI is *PI* − *NL* = 57.4. It is therefore valuable to learn about the population dynamics and per-capita growth rate, as this learning is expected to improve performance on the management objective.

Value of information analyses are typically applied for static, one-time conservation decisions, rather than sequential problems, often in the context of understanding when to act and when to delay action until after further research (Bolam et al. 2019). To better understand the value of information in the context of adaptive management, future work should investigate quantifying VoI by iteratively resolving uncertainty, requiring evaluating the gain of implementing an adaptive policy rather than a single action (Chadès et al. 2017).

#### 4.4.2 Passive adaptive management

Passive adaptive management involves learning from the outcomes of previous management actions to update knowledge about system dynamics and improve management performance over time. Decisions at each time step are optimized assuming that current knowledge of the system will not change into the future. Learning therefore occurs by monitoring the effect of an implemented action, rather than during the optimization procedure as in active adaptive management described below (Chadès et al. 2017).

Solving a passive adaptive management problem includes an implementation step and an updating step at every time point *t*, based on the current belief in model *k*, *b_t_*(*k*). The problem must be solved at every time step because the transition matrices change as the belief changes. During the implementation step, the optimal action at time *t* is selected using a weighted averaging approach, where the transition probabilities are averaged across all models with the weights given by the current belief, *b_t_*(*k*) (Williams et al. 2011). This can be solved using SDP:

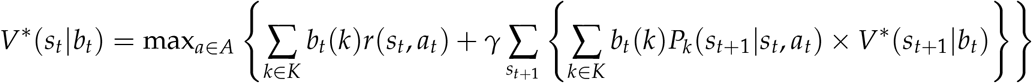

During the updating step, the resulting state, *s_t_*_+1_, after taking action *a_t_* is observed, and belief, *b_t_*_+1_, is updated according to Bayes’ rule:

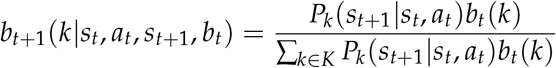

For coypu control decisions, we solve the passive adaptive management problem by updating the belief about the coypu population dynamics to improve management performance over time. We begin with an equal belief, *b_t_*_=1_(*k*) = 1/5, across the five models in the model set (Figure 5C). We iterate between 1) calculating the optimal action given the current belief state, 2) implementing the optimal action and simulating the next state, *s_t_*_+1_, given the true population dynamics, and 3) updating the belief state based observing the next state given action *a_t_*. Here we assume the state is perfectly observed. We simulate the application of passive adaptive management over 30 time steps. The belief in each model changes over time (Figure 5C), resulting in the highest belief for the two models closest to the true population dynamics *m* = 1 and *m* = 1.3 (Figure 5C). Through updating the belief state over time and implementing the optimal action given the current belief state, the management performance improves over time (Figure 5D).

#### 4.4.3 Active adaptive management

Passive adaptive management chooses the action that is optimal given current knowledge, then learns from the outcome to improve decision-making over time. In contrast, active adaptive management explicitly values future learning when selecting today’s action. Management actions are therefore chosen not only because they improve the ecological state immediately, but also because they are expected to reveal information that will improve future decisions and maximize the chance of achieving an objective over the long term.

In passive adaptive management, the SDP optimization occurs at every time step, given the current belief, and the belief state is updated as learning occurs. In active adaptive management, by contrast, the optimization only occurs once and requires that the trajectory of belief *b_t_* to *b_t_*_+1_ and state transitions be calculated for all time steps during the optimization (Chadès et al. 2017). An active adaptive policy improves management in the long term by accounting for how the belief state will change over time as a result of an action, rather than assuming a static belief state at each time step, as in passive adaptive management. The optimal value function *V*^∗^ that characterizes the performance of a policy is now a function of both the state of the system *s_t_* and belief over the models *b_t_* (i.e., *V*^∗^(*s_t_*, *b_t_*)).

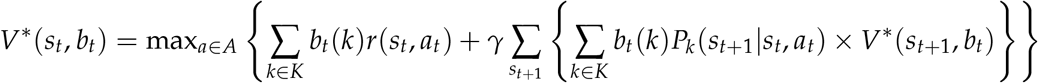

Active adaptive management with model uncertainty can be solved with many methods, including those developed for dealing with partial observability (Williams et al. 2011, Chadès et al. 2017). Here, we use a POMDP characterized by a tuple *<X*, *A*, *O*, *P*, *Z*, *r*, *γ>*, where *X* = *S* × *K* represents the factored state space with *S* denoting the possible conditions of the system and *K* denoting the model set. The unknown model is an unobserved state variable that must be inferred through observation. Similar to the partial observability section above, we can solve the POMDP by approximating the value function *V*^∗^(*s_t_*, *b_t_*) associated with the optimal policy *π*^∗^ using a set of *α*-vectors (Kurniawati et al. 2008).

For coypu control decisions, we solve the active adaptive management problem by augmenting the state space with the belief in two models where *m* ∈ {0.7, 1.3}. We choose a smaller model set than in the passive adaptive management problem due to the computational cost associated with exploring the belief and state spaces over the entire time period (i.e., curse of dimensionality). We assume a uniform belief across models and assume perfect observation of the state (i.e., *o_t_* = *s_t_*).

We also use the coypu example to demonstrate the “dual control problem” (Walters and Hilborn 1978). A management action altering a poorly understood system can produce two types of benefits: 1) short-term payoffs and 2) learning about the system so as to improve payoffs in the long-term. There is a trade-off between these two benefits, as more intensive management actions may generate information more rapidly but can also incur higher short-term costs; the challenge is to optimally balance these benefits. Here we explore the dual control problem by selecting two different discount factors, *γ* ∈ {0.95, 0.75}. The higher of which represents a decision maker who values long-term rewards more highly, and the lower of which represents a decision maker who values short-term rewards more highly.

The active adaptive management solution describes the optimal action (*a_t_*, removal effort), given the state (*s_t_*, coypu abundance), initial belief (*b_t_*_=1_, uniform across models), and discount factor. The solution differs depending on how future rewards are discounted in the optimization. At low coypu abundance, the optimal removal effort increases as the discount factor increases. With a higher discount factor, future system states are valued more highly; even though a higher removal effort is costly today, this higher removal effort will accelerate learning to yield higher rewards in the long term (Figure 5E). In contrast, with a lower discount factor, active learning becomes less useful, as the price paid for not following the passive-adaptive (optimal right now) solution exceeds the information gained from learning (Figure 5F).

## 6. Conclusions

Stochastic dynamic programming provides a powerful and flexible framework for linking ecological models directly to management decisions. Rather than focusing solely on predicting invasion dynamics, SDP identifies state-dependent management policies that explicitly account for ecological uncertainty, future consequences of present actions, and the trade-offs between management costs and ecological impacts (Bellman 1957, Williams 1982). The worked examples presented here illustrate how the same decision-theoretic framework can accommodate progressively richer sources of uncertainty, from stochastic population dynamics and spatial spread to imperfect detection and uncertainty about ecological processes themselves. Together, these approaches provide a coherent framework for making transparent, reproducible, and objective management decisions for invasive species.

Several methodological developments offer promising opportunities for extending these approaches. Future work could focus on scaling dynamic programming methods to larger spatial systems and higher-dimensional state spaces through approximate dynamic programming and reinforcement learning (Powell 2011, Bertsekas 2012). Future efforts could also be directed at improving the representation of ecological processes and observation uncertainty, and at incorporating multiple management objectives such as biodiversity conservation, ecosystem services, economic costs, and social acceptability within a unified decision framework (Gregory et al. 2012).

Although SDP remains computationally demanding for complex ecological systems, continuing advances in optimization algorithms, statistical modelling, and computing power are rapidly expanding its practical applicability (Nicol and Chadès 2011, Bertsekas 2012). Ultimately, successful invasive species management depends not only on understanding ecological systems, but also on making better decisions. By explicitly linking ecological predictions to management actions, SDP and its extensions provide a principled framework for translating ecological knowledge into effective management. We hope that the worked examples and reproducible R code presented here will encourage ecologists and practitioners to adopt decision-theoretic approaches and help bridge the gap between ecological prediction and conservation action.

## Acknowledgements

This research was supported by the ExposUM Institute of the University of Montpellier and the ANR through the project NACHOS for “Interdisciplinary approach to small carnivores - humans relationships” (grant ANR-25-CE03-5469). This research is also based upon work supported by the U.S. Department of Energy, Office of Science, Office of Advanced Scientific Computing Research, under Award Number DE-SC0024386 and the U.S. National Science Foundation under Grant No. DBI-1942280.

## Declaration of generative AI and AI-assisted technologies in the writing process

During the preparation of this work, the author used ChatGPT to polish the text and enhance the English language. After using this tool, the author reviewed and edited the content as needed and take full responsibility for the content of the published article.

## Data availability statement

Codes and data are available at https://github.com/oliviergimenez/SDPaper.

## Author contributions

O. Gimenez, A.G. Keller and C. Speakman conceived the study, developed the methodology, conducted the analyses. O. Gimenez, A.G. Keller, L. Marescot and C. Speakman wrote the original manuscript draft. All authors contributed to the interpretation of the work and critically revised the manuscript. All authors read and approved the final version of the manuscript.

